# What happened when? Brain and behavioral responses to modified structure and content in episodic cueing

**DOI:** 10.1101/2021.11.10.467871

**Authors:** Sophie Siestrup, Benjamin Jainta, Nadiya El-Sourani, Ima Trempler, Oliver T. Wolf, Sen Cheng, Ricarda I. Schubotz

## Abstract

Episodic memories are not static but can be modified on the basis of new experiences, potentially allowing us to make valid predictions in the face of an ever-changing environment. Recent research has identified prediction errors during retrieval as a possible trigger for such changes. In the present study, we used modified episodic cues to investigate whether different types of mnemonic prediction errors modulate brain activity and subsequent memory performance. Participants encoded different episodes which consisted of short toy stories. During a subsequent functional magnetic resonance imaging (fMRI) session, participants were then presented videos showing the original episodes, or modified versions thereof. In modified videos, either the order of two subsequent action steps was changed or an object was exchanged for another. Content modifications recruited inferior frontal, parietal, and temporo-occipital areas reflecting the processing of the new object information, while brain responses to structure modifications in right dorsal premotor and inferior frontal areas remained subthreshold. In a post-fMRI memory test, the participants’ tendency to accept modified episodes as originally encoded increased significantly when they had been presented modified versions already during the fMRI session. Our study provides valuable initial insights into the neural processing of different types of episodic prediction errors and their influence on subsequent memory.

## 1 INTRODUCTION

Episodic memories enable us to vividly re-live events that we experienced at some point in our personal life (Tulving, 2002). But there is evidence that they are not always veridical reconstructions of our past. On the contrary, memories are flexible and prone to change (Nader & Einarsson, 2010; Nader, 2015; Lee et al., 2017; Scully et al., 2017). In everyday life, situations we encounter are usually not exactly the same as those we experienced before. So, there is always a certain discrepancy between our expectations, which we derive from our memories, and the events we then actually experience. According to the predictive coding framework, this discrepancy leads to a prediction error (Barto et al., 2013; Reichardt et al., 2020). Generally, prediction errors serve as bottom-up learning signals that allow us to adapt our internal predictive models to an ever-changing environment, so that we can maintain valid predictions in the long run (Friston, 2005; Friston & Kiebel, 2009; Schubotz, 2015). Accordingly, it is likely adaptive that memories are modified in favor of valid internal models informed and updated by later experiences (Exton-McGuinness et al., 2015; Fernández et al., 2016). Indeed, recent research has identified mnemonic prediction errors as potential drivers of memory change (Sinclair & Barense, 2019).

In the present study, we examined the neural and behavioral level of mnemonic prediction errors during episodic retrieval by targeting two basic types of episodic memory information: either their content (“what”) or their structure (“when”) (cf. Griffiths et al., 1999). To do so, we employed a new episodic cueing paradigm based on previous work (Schiffer et al., 2012; 2013). After encoding short episodes from videos, participants went through a functional magnetic resonance imaging (fMRI) session and were either presented original episode videos or slightly modified versions thereof. A subset of videos was manipulated with regard to the occurrence of an object (content modification) or the order of two consecutive action steps (structure modification) to elicit different types of mnemonic prediction errors. In a post-fMRI memory test, participants’ memory for original and modified episodes was probed.

If new content and/or structure information elicited changes in the predictive model during fMRI, we expected that this would reduce memory performance in the post-fMRI memory test (Schiffer et al., 2012; 2013). This could be reflected in a weakening of the original episodic memory, i.e., false rejection of original videos in the memory test, and/or in the creation of alternative episode representations, i.e., false acceptance of modified videos in the memory test.

Concerning brain activations, we expected that some regions would respond to both types of episodic modifications. Particularly, the medial frontal cortex (MFC) is a candidate area that may serve more general control over consolidation and retrieval of long-term memories (Euston et al., 2012; Peters, et al., 2013). Furthermore, the hippocampus is regarded a core structure of episodic memory (Maguire et al., 2016; Horner & Doeller, 2017; Stachenfeld et al., 2017) and responds to mnemonic prediction errors (Long et al., 2016; Bein et al., 2020). Further, we expected increased activity in brain areas related to action observation (Caspers et al., 2010) since we previously found them attenuated when subjects adapted to episode modifications affecting both objects and sequential structure combined (Schiffer et al., 2012; 2013).

Apart from these common effects, structure and content episodic modifications were also expected to affect different regions. Structure modifications should elevate activity in premotor areas due to their central role in sequential order processing (Schubotz, 2004). Dorsal premotor and adjacent prefrontal sites along the superior frontal sulcus (PMd/SFS) were found for stepwise ordinal linking of individual action steps, as required in quite different, but especially predictive, tasks (Stadler et al., 2011; Schubotz et al., 2012; Tamber-Rosenau et al., 2011; Hrka et al., 2014; Pomp et al., 2021). By contrast, content modifications were expected to engage areas re^ć^lated to object processing, including lateral occipitotemporal cortex (OTC) (Lingnau & Downing, 2015), anterior intraparietal sulcus (IPS) (Creem-Regehr, 2009; Schubotz et al., 2014), and fusiform gyrus (FG; Reber et al., 2005). Finally, based on earlier findings, we tested the hypothesis that within the inferior frontal gyrus (IFG), Brodmann area 44 (BA 44) may be especially involved in structure modification due to its role in processing the sequential structure of actions (Fiebach & Schubotz, 2006; Clerget et al., 2009) whereas BA 45 may be more relevant for semantic processing, hence also content modification of episodes (Badre & Wagner, 2007; Heim et al., 2009).

## 2 MATERIALS AND METHODS

### 2.1 **Participants**

Forty female participants took part in the study. Only females participated in order to achieve a good match between participants’ hands and the actress’s hands in the stimulus material, as participants were supposed to believe they were shown in the videos themselves. Two participants did not return for the fMRI session after training. Additionally, data from one participant had to be excluded from analysis due to nausea and dizziness during the fMRI session. In another case, an unknown technical problem led to the incomplete acquisition of functional data. Thus, the final sample for fMRI analyses consisted of 36 participants, age 18 – 28 years (*M* = 22.67 years, *SD* = 2.40 years). Participants of the final sample were all right-handed as assessed by the Edinburgh Handedness Inventory (Oldfield, 1971). Handedness scores varied from +60 to +100 (*M* = 92.17, *SD* = 10.95). Data from one participant was not used for the analysis of the post-fMRI memory test due to misunderstood instructions. All participants had normal or corrected-to-normal vision, were native German speakers, and reported no history of neurological or psychiatric disorders or substance abuse. Participants signed a written informed consent and received monetary compensation or course credits for their participation. The study was conducted in accordance with the Declaration of Helsinki and approved by the Local Ethics Committee of the University of Münster.

### 2.2 Stimuli

Even though episodic memory can potentially never be completely independent of semantic knowledge (Cheng et al., 2016) we wanted to reduce influences of everyday experiences as far as possible in our study. To this end we chose to use abstract but complex toy-based stories so that we could ensure controlled encoding of unique episodes in the laboratory.

Stimuli were 78 short videos (duration = 8.80 – 17.88 s, *M* = 12.71 s) of stories that were played with PLAYMOBIL® toys, showing only the toys and hands and underarms of an actress. Stories comprised six to nine action steps (*M* = 7.4 steps) and four to 14 separable objects (*M* = 6.93 objects), such as characters, animals, vehicles and tools. The exact same object appeared in only one of the stories.

Stories were filmed from above with a digital single-lens reflex camera (Nikon D5300) which was centrally mounted above the table and faced straight down. Matt white paper served as a base. A frame of 47.5 cm x 28 cm was taped on the paper, congruent with the section captured by the camera (in the following referred to as camera frame). Objects that were needed for a particular story were positioned next to the camera frame and were only moved into view in the moment at which they appeared in the story. During filming, the actress wore a black pullover and black rubber gloves. To facilitate future imitation from demo videos, the back of the right hand was marked with a yellow dot (Franz et al., 2007). Video material was edited using Adobe Premiere Pro CC (Adobe Systems Software, Dublin, Ireland, Version 12.1.2). All videos had a frame of size 1920 x 1080 pixels and a frame rate of 25 frames per second. Videos started with seven frames showing only background and ended after seven frames showing the final toy constellation. The original perspective of the videos was the third-person (observer) perspective (3pp) and they were rotated 180° to create their first-person (field) perspective (1pp) counterparts using Adobe Premiere Pro CC (Wurm & Schubotz, 2018). Throughout the experiment, videos were presented at a visual angle of approximately 7.3° x 13° using the stimulus presentation software Presentation (version 20.3 02.01.19, NeuroBehavioral Systems, Berkeley, CA, USA).

On the basis of two pilot studies, we chose 24 out of originally 30 stories for our stimulus set. Stories were excluded when they were particularly difficult to imitate or describe. One of the 30 stories was excluded due to low memorability as indicated by low performance in a signal detection task.

The 24 final stories existed in three different versions each: (1) an original version (*ori*) as encoded by the participants, (2) a version in which two adjacent action steps were switched (structure modification) (*str*) and (3) another variation of the original video in which one object was exchanged (content modification) (*con*). Story scripts were created by five experimenters who all had to agree that the original story as well as modifications thereof were semantically valid (within a toy world) and that modifications did not change the overall outcome of the story. For creating videos with modifications, the respective stories were played and filmed again exactly the way as for the original video. The only aspect that differed between original and modified versions was a single change of either the order of two action steps (i.e., one transition out of 7.33 transitions on average, for structure modifications) or one object (i.e., one object out of 6.95 objects on average, for content modifications). Modifications were never introduced in the first two action steps so that the beginning of a video served as a cue for prediction. Furthermore, no modifications were introduced in the last two action steps either. The exact time point of the modification in each video was determined by identifying the video frame which diverged from the original version. Six other stories were used in one version only. Four of them were presented for the first time in the fMRI session, we refer to them as untrained episodes in the following. The two remaining videos were only used for practice and did not appear in the fMRI experiment.

For examples of different story versions, see Figure 1 and Supplementary Table 1. Videos are available upon reasonable request via the Action Video Corpus Muenster (https://www.uni-muenster.de/IVV5PSY/AvicomSrv/).

**Figure 1.**
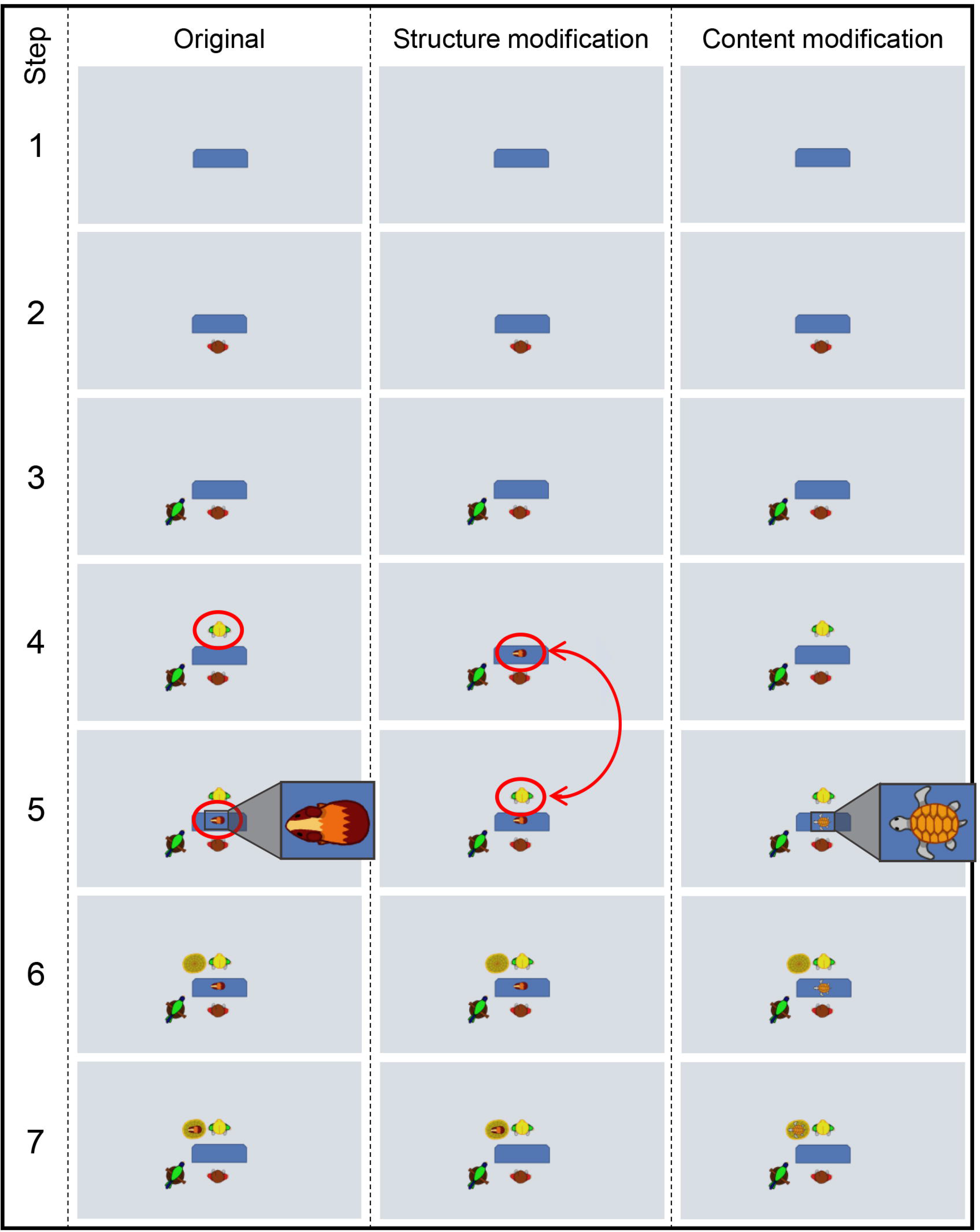
Example of an original episode and its modified versions, shown by the sequence of the main event steps. Twenty-four stories existed in three different versions each: an original, a structure modification, and a content modification. For the structure modifications, two adjacent action steps were switched compared to the original. In this example, the original shows the blonde woman join the scene before the guinea pig is positioned on the sales counter; in the structure modification, the guinea pig appears before the blonde woman (red circles). For the content modifications, an object was exchanged compared to the original (here: tortoise instead of guinea pig on the sales counter in step 5). Note that in the fMRI experiment, each participant was only presented with one of the three versions of a story. We do not reproduce photos of our stimulus material because it is copyrighted material (PLAYMOBIL® figures); instead, we provide schematic images.

### 2.3 Procedure

#### 2.3.1 **Training**

Training sessions were conducted in a computer lab at the Department of Psychology at the University of Münster. The training consisted of two sessions that took place on two consecutive days and lasted about 2 h and 1.5 h, respectively. During each of the two sessions, participants encoded half of the episodes. We chose to split the training over two days to avoid fatigue or a decrease in motivation due to the relatively long duration of the task. For encoding episodes, participants imitated half of the 24 stories from demo videos (i.e., the original versions) and only observed the other half, equally distributed over both training days (factor AGENCY).

The 24 demo videos were organized in four subsets, containing six videos each, balanced for the number of action steps (*A1*, *A2*, *B1*, *B2*). Participants either imitated videos belonging to *A* subsets and observed videos belonging to *B* subsets or vice versa. In each training session, participants went through one imitation block and one observation block, each consisting of one subset of videos. This means that each participant trained each video either during session one or during session two, the same video was not trained on both days. The order of videos within these blocks was randomized for each participant. All stories were imitated and only observed equally often over the course of the study. The first training session started with a short practice phase of the task (2 videos). Then, half of the participants started with imitation, the other half with observation. The same applied to the second training session, except that participants who had started with imitation on the first day now started with observation, and vice versa. The order of blocks and the order in which video subsets were trained was balanced between participants. Participants always went through each story eight times, including four times in 3pp and another four times in 1pp. If the story was to be imitated, participants first watched the story four times in 3pp, then once in 1pp and then performed the last three repetitions of the story themselves. If the story was only observed, the video was instead presented three more times in 1pp. This way, participants experienced each story equally often from each visual perspective, allowing us to establish the factor PERSPECTIVE in the fMRI session through the presentation of just one of both perspectives. The optimal number of video presentations for successful imitation had previously been determined in a pilot study.

During training, the participant sat at the same setup which had been used for filming the stimulus material and likewise wore a black pullover and gloves with a yellow dot on the right hand. The experimenter sat opposite of the participant, supervising the performance (Figure 2). For each story, the toys were positioned next to the camera frame, following the same arrangement as used while creating the stimulus material. After imitation or observation, participants had to deliver a detailed description of the story to ensure that they understood it correctly and had paid attention to all objects involved. If participants made a mistake during an imitation or description trial, they were immediately interrupted by the experimenter to avoid encoding of incorrect scripts. They would then start over with a new imitation/description attempt.

**Figure 2.**
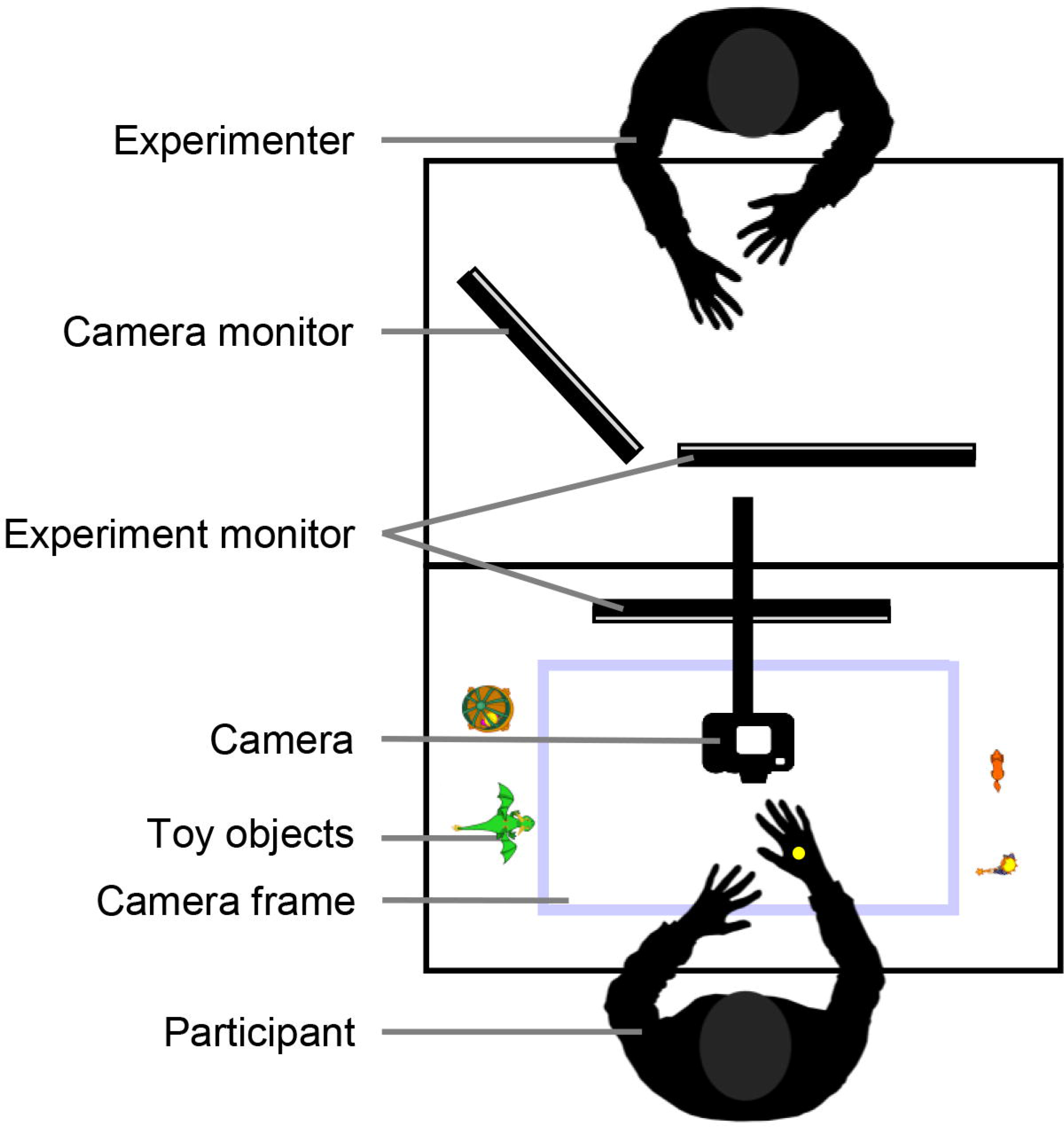
Training setup. During training participants observed and imitated toy stories, while sitting at the filming setup. Their performance was monitored by the experimenter.

The two described experimental factors AGENCY (*imitation*, *observation*) and PERSPECTIVE (*1pp*, *3pp*) are not further addressed in the present paper, as they are central for a companion paper (Jainta et al., 2021) describing the influence of personal involvement on recall. Both factors were balanced with respect to the factors reported here, so we can exclude any confounding effects (fully-crossed design).

#### 2.3.2 fMRI session

The fMRI session took place approximately one week after the second training session (range = 7 - 12 days; *M* = 7.42 days, *SD* = 0.90 days). Participants were told that videos of themselves playing the stories would be presented in the fMRI session. Even though participants had actually been filmed during training, these videos were not used during the fMRI experiment. This was only a cover story to elevate personal identification with the videos to benefit episode reactivation. Participants were fully debriefed after completion of the study.

During the fMRI session, participants were presented with original and modified videos reminiscent of the previously encoded episodes. Importantly, each video was only shown in the original or one divergent version. Following a previously used paradigm (Schiffer et al., 2012; 2013), modified and original episodes were presented repeatedly to simulate the natural circumstances that potentially foster memory modification, i.e. updating of internal models due to increasing evidence for the validity of an alternative. Thus, eight videos were repeatedly presented in the original version, eight included a structure modification and eight a content modification. Which stories belonged to which conditions was counterbalanced between participants. Half of the videos were shown in 1pp, the other half in 3pp, perfectly balanced over conditions. Additionally, four untrained stories were included in the fMRI session.

The fMRI experiment consisted of six blocks, each containing the 24 videos reminiscent of the previously encoded episodes. Consequently, each video was presented six times over the course of the session. Within blocks, videos were presented in pseudo-random order so that transition probabilities between conditions were balanced. Additionally, each block contained three null events during which only a fixation cross was presented (duration: 7 – 10 s). Furthermore, each untrained video was presented once per block. Therefore, the whole experiment contained 18 null events and 24 untrained video trials. Participants were not informed about the block-structure of the experiment.

Participants were instructed to attentively watch the presented videos. They were told that after some videos a short description would be presented (e.g. “Rescuing princess”) that either matched or did not match the story shown in the video (question trials). The task was to either accept or reject the description by pressing one of two buttons on a response box with the right index or middle finger, respectively. This type of task has been used successfully before to focus participants’ attention on complex video stimuli (El-Sourani et al., 2019). The descriptions represented the stories on a gist-level, so specific details were irrelevant to the correct answer (Figure 3). Throughout the entire experiment, each story was once followed by a matching description and once by a non-matching description, resulting in a total number of 56 question trials in the experiment. Each block contained nine to ten question trials and per block, approximately 50 % of descriptions were to be accepted, 50 % were to be rejected. The question was presented for a maximum of 3 s or until participants responded. Upon response delivery, participants received 1 s written feedback whether they answered correctly, incorrectly, or too late, in case no response was given. Participants were naïve with regard to this distribution of question trials.

**Figure 3.**
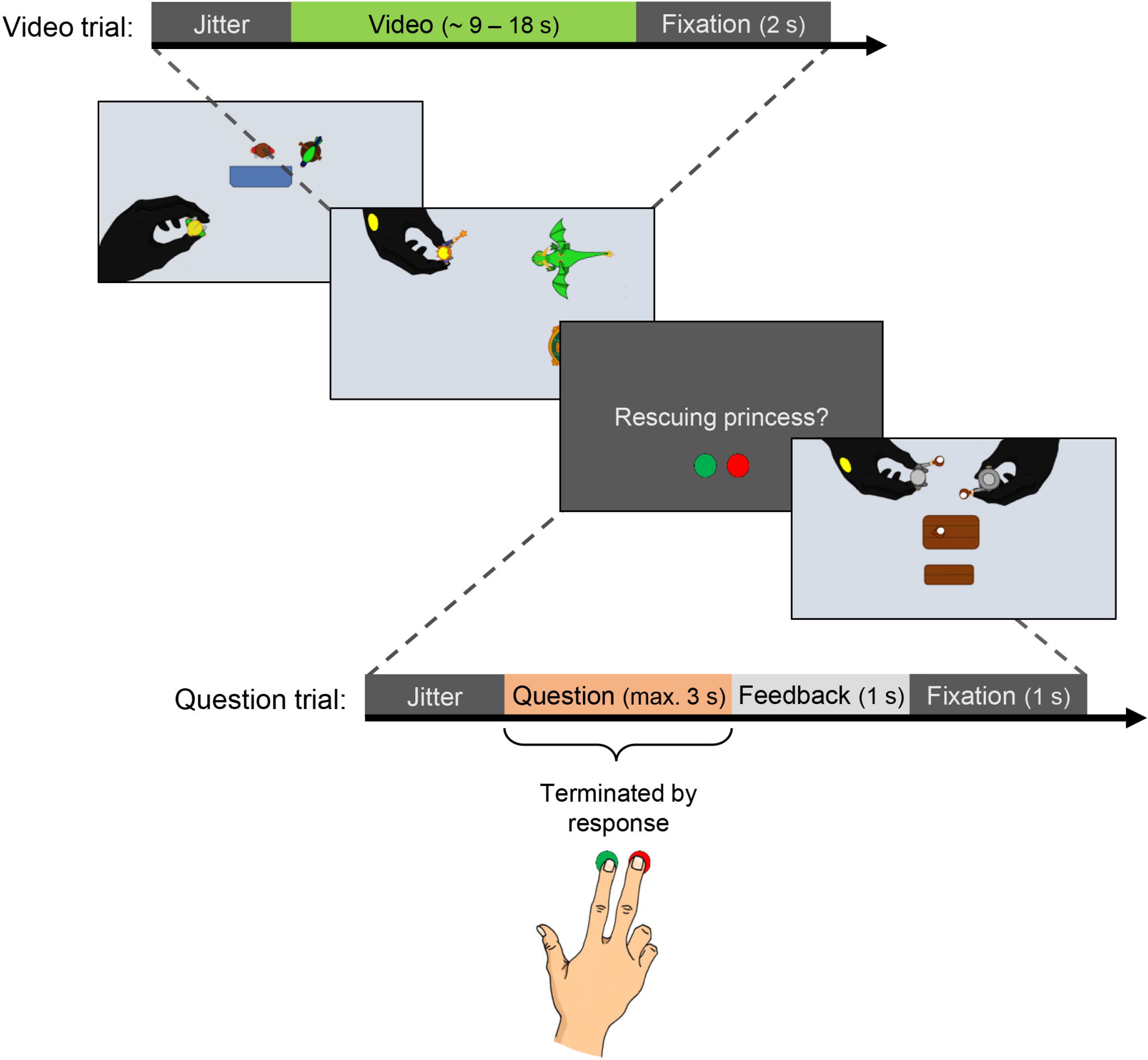
Schematic depiction of task during fMRI session. Video trials consisted of a variable jitter (0, 0.5, 1, or 1.5 s of fixation), a video showing a toy story (ca. 9 – 18 s) and a 2 s interstimulus interval (fixation). Question trials included a variable jitter, a question regarding the story shown in the preceding video (maximally 3 s long or terminated by response), and a 2 s interstimulus interval. The interstimulus interval after question trials was divided into 1 s feedback (“correct”, “incorrect”, “too late”) and 1 s fixation. Aside from the question, it was depicted which button should be pressed to accept (left, green) or reject (right, red) the description.

Between trials, a fixation cross was presented for a duration of 2 s (1 s after question trials) to serve as an interstimulus interval. Before each trial, a variable jitter of 0, 0.5, 1 or 1.5 s of fixation was added for enhancement of the temporal resolution of the BOLD response (Figure 3).

In total, the fMRI task had a duration of approximately 48 min. The task had previously been practiced at the end of the second training session (four video trials, four question trials, one null event). Practice videos were shown twice each and not re-used in the fMRI session.

#### 2.3.3 Post-fMRI memory test

Immediately after the fMRI session, participants completed an explicit memory test. Importantly, encoding occurred incidentally, as participants were not informed beforehand that their memory for episodes would be tested.

Participants were seated in a separate room in front of a laptop and instructed to remember their training sessions one week prior. They were presented all stories that they had seen in the fMRI session in two different versions. More precisely, when modified videos had been presented during the fMRI session, these modified videos were presented again during the memory test and *additionally* each story was shown in the original version. When original episode videos had been presented during the fMRI scan, these original videos were presented again in the memory test and *additionally*, each story was shown in a modified version, *either* containing a structure modification in half of the cases *or* a content modification.

The participants’ task was to rate after each video whether they knew this exact episode from the training, using a Likert scale including 1 (yes), 2 (rather yes), 3 (rather no) and 4 (no), by pressing one out of four marked keys on the laptop’s keyboard. Response time was not restricted but participants were instructed to respond quickly and intuitively. Videos were presented in a pseudo-randomized order, so that half of the stories (of each experimental condition) were first presented in their original version followed by a modified version and vice versa. Untrained videos were shown twice in the same version, so that, in total, the memory test comprised 56 video trials. The completion of the task took approximately 15 min.

### 2.4 MRI data acquisition and preprocessing

MRI scans were conducted with a 3-Tesla Siemens Magnetom Prisma MR tomograph (Siemens, Erlangen, Germany) using a 20-channel head coil. Participants lay supine on the scanner bed with their right index and middle finger positioned on the two appropriate buttons on a response box. Head, arm and hand movements were minimized by tight fixation with form-fitting cushions. Participants were provided with earplugs and headphones to attenuate scanner noise. Stimuli were projected on a screen that the participants saw via an individually adjusted mirror which was mounted on the head coil.

High resolution T1 weighted anatomical images were obtained with a 3D-multiplanar rapidly acquired gradient-echo (MPRAGE) sequence prior to functional imaging. 192 slices with a thickness of 1 mm were acquired, using a repetition time (TR) of 2130 ms, an echo time (TE) of 2.28 ms, a flip angle of 8° and a field of view (FoV) of 256 x 256 mm². Functional images of the whole brain were acquired in interleaved order along the AC–PC plane using a gradient-echo echoplanar imaging (EPI) sequence to measure blood-oxygen-level-dependent (BOLD) contrast. Thirty-three axial slices with a thickness of 3 mm were obtained in an interleaved order, using a TR of 2000 ms, a TE of 30 ms, a FoV of 192 x 192 mm² and a flip angle of 90°.

Processing of imaging data was conducted with SPM12 (Wellcome Trust, London, England) implemented in MATLAB (version R2018b, The MathWorks Inc., Natick, MA, USA). Data were preprocessed by slice time correction to the middle slice, movement correction and realignment to the mean image, co-registration of the functional data to individual structural scans, normalization of functional and structural images into the standard MNI space (Montreal Neurological Institute, Montreal, QC, Canada) on the basis of segmentation parameters and spatial smoothing using a Gaussian kernel of full-width at half maximum (FWHM) of 8 mm. Further, a 128 s high-pass temporal filter was applied.

### 2.5 Statistical data analysis

#### 2.5.1 **Behavioral data**

The behavioral data analysis was conducted using RStudio (R Core Team, 2020; version 1.3.1073).

To check our hypothesis that repeated presentation of modified videos in the fMRI session leads to a decrease in memory performance in the memory test, we considered the discrimination index *P_r_* (Snodgrass & Corwin, 1988). In addition, we evaluated ratings for modified and unmodified videos separately to assess the extent of erroneous acceptance of modified videos as originally encoded and/or erroneous rejection of original episodes. Further, we examined reaction times in the memory test, which can serve as an indicator of how long it takes to retrieve information (correctly) from memory (Collins & Quillian, 1969). Longer reaction times indicate increased difficulty of retrieval due to higher cognitive processing demands (Larsen & Plunkett, 1987; Noppeney & Price, 2004), which may also occur when competing versions of an episode must be processed.

For the analysis of *P_r_* values (unmodified videos = targets, modified videos = distractors) as well as ratings and reaction times for modified videos in the memory test (modified_MT_), we applied a 2 x 2 within-subject factorial design with the factors MODIFICATION_FMRI_ (*yes, no*) and VERSION_MT_ (*str, con*). For analyzing responses to original videos in the memory test (original_MT_), we applied a within- subject design with the factor VERSION_FMRI_ (*ori*, *str*, *con*).

For each factorial combination, we calculated the discrimination index *P_r_* (hit rate minus false alarm rate; ratings “yes” and “rather yes” were grouped as “acceptance”, and “no” and “rather no” as “rejection”), the average rating, and the average reaction time for correct responses. We report additional analyses on hit rates and false alarm rates in the Supplementary Material. Finally, we compared ratings for untrained video to those for original_MT_ and modified_MT_ videos. Please note that low ratings indicate acceptance, while high ratings indicate rejection.

We also conducted an explorative analysis on behavioral data from the fMRI session. We calculated the error rate and mean reaction time according to the within-subject factor VESRION_FMRI_ (*ori*, *str*, *con*) per participant. No response was given in only 0.7 % of all question trials, and these trials were not further considered in the analysis.

For the choice of statistical tests, data were inspected for normal distribution using the Shapiro Wilk Test. Further, data were checked for extreme outliers as defined as values above quartile 3 + 3* interquartile range (*IQR*) or lower than quartile 1 – 3 * *IQR*. When data were normally distributed or could be transformed to fit normal distribution (reaction times for original videos, reaction times during fMRI task; logarithmic transformation) and showed no extreme outliers, we used repeated measures analyses of variance (rmANOVA). Post-hoc pair-wise comparisons were conducted with paired *t*-tests (one-tailed when comparing *ori* and *str* and *ori* and *con*, two-tailed when comparing *str* and *con*). When the prerequisites for parametric analysis were not met, we used a non-parametric rmANOVA based on aligned rank data (package *ARTool*; Wobbrock et al., 2011; *P_r_* values, ratings, error rates). Several participants did not give any correct answers (i.e. rejection) in response to modified_MT_ videos for one or more factorial combinations. To avoid the listwise exclusion of many datapoints for reaction times when using a rmANOVA approach, we decided to use one-tailed Wilcoxon signed rank tests when comparing reaction time between levels of the factor MODIFICATION_FMRI_, separately for each modification type. This way, we could include 16 datapoints for structure and 29 datapoints for content modified videos. We additionally excluded another datapoint for content modifications due to a very long mean reaction time in the *yes* condition (8.85 s) which likely does not represent purely mnemonic processes. Please note that this did not influence the overall outcome of the analysis.

When parametric statistical approaches were used, we report (untransformed) mean values and standard errors of the mean as descriptive statistics, when non-parametric approaches were applied, we report median values and *IQR*. For all behavioral analyses, we applied a significance level of α = 0.05. *P*-values were adjusted according to the Bonferroni correction for multiple comparisons (Bonferroni, 1936).

#### 2.5.2 fMRI design specifications

Statistical analyses of the fMRI data were conducted with SPM12. We used a general linear model (GLM) for serially autocorrelated observations (Friston et al., 1994; Worsley & Friston, 1995) and convolved regressors with the canonical hemodynamic response function. Regressors were original videos (*ori*), videos containing a structure modification (*str*) and videos containing a content modification (*con*), each comprising 48 trials. For *str* and *con* trials, the onsets of events were time-locked to the point in the video at which the modification occurred (time of modification). For *ori* trials, we calculated a hypothetical time of modification (mean of times that corresponded to points of structure and content modification in the non-modified video) to serve as a comparable onset. These conditions were modeled as events as we were interested in the phasic effect of the prediction violation at the precise moment it occurred. To each of those regressors, we added a parametric modulator to model the repeated presentation of each video. The 24 untrained videos were modelled as events as well, with onsets timed to the middle of the video. Two additional regressors modeled the 18 null events and the 56 question trials. The modeled activation of null events and questions was time-locked to their respective onsets. Null events were modeled as epochs, containing their full presentation time (7 – 10 s), while questions were modeled as events. The six subject-specific rigid-body transformations obtained from realignment were included as regressors of no interest. Therefore, the GLM comprised 15 regressors in total.

We calculated first-level-*t*-contrasts for *str* > *ori* and *con* > *ori* to analyze brain activity in response to the specific modification types. Grey matter masking was applied on the first level of the analysis. For masking, we used the smoothed individual normalized grey matter image (8 mm FWHM) which was thresholded at 0.2 using ImCalc in SPM12 to create a binary mask. Second-level group analyses were performed by using one-sample *t*-tests across participants. A conjunction of *str* > *ori* and *con* > *ori* contrasts was calculated to detect shared effects of both modifications (Nichols et al., 2005). We applied false discovery rate (FDR) correction and a threshold of *p* < 0.05 (peak level). In case no significant activation was detected using this threshold, we applied no correction for multiple comparison with *p* < 0.001. Brain activation patterns were visualized with the software MRIcroGL (version 1.2.20200331, McCausland Center for Brain Imaging, University of South Carolina, USA).

To test our hypothesis that BA 44 (IFG pars opercularis) is more involved in detecting structure modification than content changes in episodes, while the reverse pattern may hold for BA 45 (IFG pars triangularis), separate structural ROIs were generated for the two areas in each hemisphere using the automated anatomical labeling atlas (Tzourio-Mazoyer et al., 2002) with the WFU PickAtlas toolbox (Maldjian et al., 2003) in SPM12. Mean beta values for the *str* > *ori* and *con* > *ori* contrasts were extracted from each ROI using the MarsBaR toolbox (Brett et al., 2002). The conditions for parametric analysis were confirmed as described above for behavioral data. We performed a rmANOVA with the factors HEMISPHERE (*left*, *right*), ROI (*BA 44*, *BA 45*), and CONTRAST (*str > ori*, *con > ori*). Post-hoc comparisons were conducted with one-tailed paired *t*-tests and we report adjusted *p*-values (Bonferroni, 1936).

## 3 RESULTS

### 3.1 **Post-fMRI memory test**

#### 3.1.1 Discrimination index *P_r_*

First, we investigated the general memory performance in the memory test, using the discrimination index *P_r_*. A non-parametric rmANOVA with the factors MODIFICATION_FMRI_ (*yes, no*) and VERSION_MT_ (*str, con*) revealed a significant main effect of MODIFICATION_FMRI_ (*F*_(1,34)_ = 20.220, *p* < 0.001, η*p²* = 0.373) which was driven by higher *P_r_* values for *no* (*Median_no_* = 0.50, *IQR_no_* = 0.375) compared to *yes* (*Median_yes_* = 0.250, *IQR_yes_* = 0.313), indicating a better memory performance when no modifications had been presented during the fMRI session. There was also a significant main effect of VERSION_MT_ (*F*_(1,34)_ = 85.828, *p* < 0.001, η*p²* = 0.716), which was explained by higher *P_r_* values for *con* (*Median_con_* = 0.688, *IQR_con_* = 0.313) than for *str* (*Median_str_* = 0.125, *IQR_str_* = 0.250). There was no significant interaction of MODIFICATION_FMRI_ and VERSION_MT_ (*F*_(1,34)_ = 2.074, *p* = 0.159, η*p²* = 0.058; Figure 4A).

**Figure 4:**
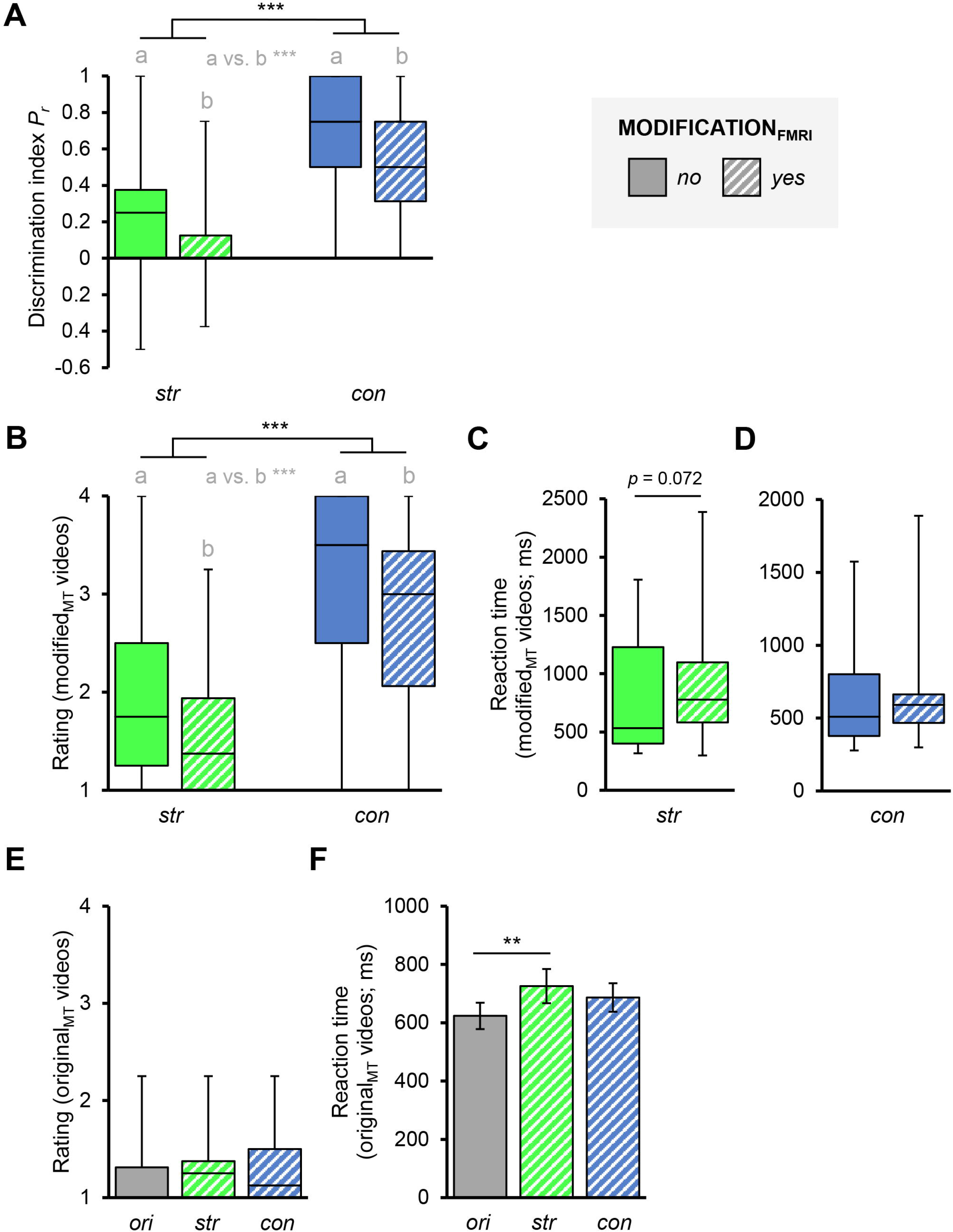
Behavioral results from post-fMRI memory test. For modified_MT_ and original_MT_ videos, participants rated whether it showed an originally encoded episode on a Likert scale including 1 (yes), 2 (rather yes), 3 (rather no) and 4 (no). **(A)** Discrimination index *P_r_* from post-fMRI memory test. Original_MT_ videos were the targets while modified_MT_ videos were distractors. Statistics: non- parametric rmANOVA with the factors MODIFICATION_FMRI_ (*yes, no*) and VERSION_MT_ (*str, con*), *n* = 35. **(B)** Ratings for modified_MT_ videos. Statistics: non-parametric rmANOVA based on aligned rank data with the factors MODIFICATION_FMRI_ (*yes, no*) and VERSION_MT_ (*str, con*), *n* = 35. **(C)** Reaction times for structure modified videos, *n* = 16. Statistics: Wilcoxon signed rank tests. **(D)** Reaction times for content modified videos, *n* = 28. **(E)** Ratings for original_MT_ videos. Statistics: non- parametric rmANOVA with the factor VERSION_FMRT_ (*ori*, *str*, *con*), *n* = 35. **(F)** Reaction times for original_MT_ videos. Statistics: rmANOVA with the factor VERSION_FMRT_ (*ori*, *str*, *con*), *n* = 35, post- hoc pairwise comparisons: paired *t*-tests, *p*-values were corrected for multiple comparisons. Box plots show medians, quartiles, minima and maxima. Bar plots show means and standard errors. ** = *p* < 0.01, *** = p < 0.001. *ori* = original, *str* = structure modification, *con* = content modification. a vs. b indicates the main effect of MODIFICATION_FMRI_.

#### 3.1.2 Ratings and reaction times for modified_MT_ videos

Next, we investigated whether the repeated presentation of modified episodes leads to the false acceptance of the same as originally encoded in the memory test. We computed a non-parametric rmANOVA with the factors MODIFICATION_FMRI_ (*yes, no*) and VERSION_MT_ (*str, con*), which revealed a significant main effect of MODIFICATION_FMRI_ (*F*_(1,34)_ = 25.554, *p* < 0.001, η*p²* = 0.429). The effect was driven by more rejective ratings for *no* (*Median_no_* = 2.625, *IQR_no_* = 0.938) than for *yes* (*Median_yes_* = 2.25, *IQR_yes_* = 1.094). Thus, when modified episodes had been presented during the fMRI session, participants were more likely to accept the same modified versions in the memory test than when original episodes had been presented. Additionally, there was a significant main effect of VERSION_MT_ (*F*_(1,34)_ = 122.647, *p* < 0.001, η*p²* = 0.783). More rejective ratings were detected for *con* (*Median_con_* = 3.250, *IQR_con_* = 1.406) than for *str* (*Median_con_* = 1.563, *IQR_con_* = 0.719), which means that participants were generally more prone to accept structurally modified videos than those with modified content. There was no significant interaction of the two factors MODIFICATION_FMRI_ and VERSION_MT_ (*F*_(1,34)_ = 2.010, *p* = 0.165, η*p²* = 0.056; Figure 4B). The same results were obtained when analyzing false alarm rates, which were significantly increased after the presentation of modified episodes in the fMRI session and, in addition, for structurally modified videos in general (cf. Supplementary Results for details).

We found a near-significant trend that reaction times were longer for structurally modified videos when this version had already been presented in the fMRI session (*Z* = -1.80, *p* = 0.072; Figure 4C). No significant difference was found concerning reaction times for videos with content modification (*Z* = -1.089, *p* = 0.276; Figure 4D).

#### 3.1.3 Ratings and reaction times for original_MT_ videos

To see whether the experience during the fMRI session influenced acceptance of original videos in the memory test, we calculated a non-parametric rmANOVA for the factor VERSION_FMRI_ (*ori*, *str*, *con*). Descriptively, ratings indicated a slightly lower acceptance of original_MT_ videos when modified videos had been presented during the fMRI session (*Median_ori_* = 1.0, *IQR_ori_* = 0.313; *Median_str_* = 1.250, *IQR_str_* = 0.375; *Median_con_* = 1.125, *IQR_con_* = 0.50), but this effect was not significant (*F*_(2,68)_ = 1.5822.010, *p* = 0.213, η*p²* = 0.044; Figure 4E). This finding was also reflected by ceiling hit rates (cf. Supplementary Results for details).

Further, a rmANOVA revealed a significant main effect of VERSION_FMRI_ on reaction times for original_MT_ videos (*F*_(2,68)_ = 3.725, *p* = 0.029, η*p²* = 0.099). Pairwise comparisons revealed that reaction times were significantly longer when structurally modified videos had been presented in the scanner compared to original videos (*t*_(34)_ = 3.287, *p* = 0.004). No significant differences were detected between the levels *con* and *ori* (*t*_(34)_ = 1.779, *p* = 0.126) and *str* and *con* (*t*_(34)_ = 0.725, *p* = 1; Figure 4F).

#### 3.1.4 Ratings for untrained_MT_ videos

To control for a general acceptance bias, we considered ratings for untrained_MT_ videos as well. A non-parametric rmANOVA on ratings for original_MT_, modified_MT_ and untrained_MT_ videos revealed a significant difference between the three types of videos (*F*_(2,68)_ = 418.16, *p* < 0.001, η*p²* = 0.925), as ratings for untrained_MT_ videos (*Median_untr_*= 4, *IQR_untr_*= 0) were significantly more rejective than for original_MT_ (*Median_ori_* = 1.208, *IQR_or i_*= 0.250; *Z* = -5.283, *p* < 0.001 ) and modified_MT_ videos (*Median_mod_* = 2.469, *IQR_mod_* = 1.031; *Z* = -5.280, *p* < 0.001). This finding indicates that participants correctly remembered that untrained videos had not been part of the training, even though they had been repeatedly presented during the fMRI session.

### 3.2 Behavioral performance during fMRI session

We calculated a non-parametric rmANOVA on error rates for the fMRI task with the factor VESRION_FMRI_ (*ori*, *str*, *con*). Descriptively, error rates were generally very low for all factor levels (*Median_ori_* = 0, *IQR_ori_* = 0; *Median_con_* = 0, *IQR_con_* = 0.02; *Median_str_* = 0, *IQR_str_* = 0) and did not differ significantly (*F*_(2,70)_ = 0.113, *p* = 0.894, η*p²* = 0.003). We also did not find a significant main effect of VESRION_FMRI_ on reaction times during the fMRI session (*F*_(2,70)_ = 0.983, *p* = 0.380, η*p²* = 0.027). These findings indicate that the participants’ attention to (and understanding of) the stories was very good and not deteriorated by the switch of single action steps or objects. Importantly, this shows that any (neural) differences between these conditions are not simply based on confusion or generally elevated cognitive effort in respect to modifications.

### 3.3 fMRI results

#### 3.3.1 **Whole brain analysis**

First, we tested whether structure and content modifications elicit common brain activation patterns compared to original episodes. The conjunction of the whole brain contrasts *str* > *ori* and *con* > *ori* did not yield significant activation after correction for multiple comparisons. Without correction and at a threshold of *p* < 0.001, there was significant activation in the opercular part of the right IFG which corresponded to BA 44 (*x* = 51, *y* = 17, *z* = 23; 48 voxels).

To identify which brain regions were active for structure modifications, we calculated the whole brain contrast *str* > *ori*. This contrast did not yield significant activation after correction for multiple comparisons. Without correction and at a threshold of *p* < 0.001, activation clusters were detected in the right hemisphere in SFS, PMd and the opercular part of the IFG (BA 44) (Figure 5 and Table 1).

**Figure 5.**
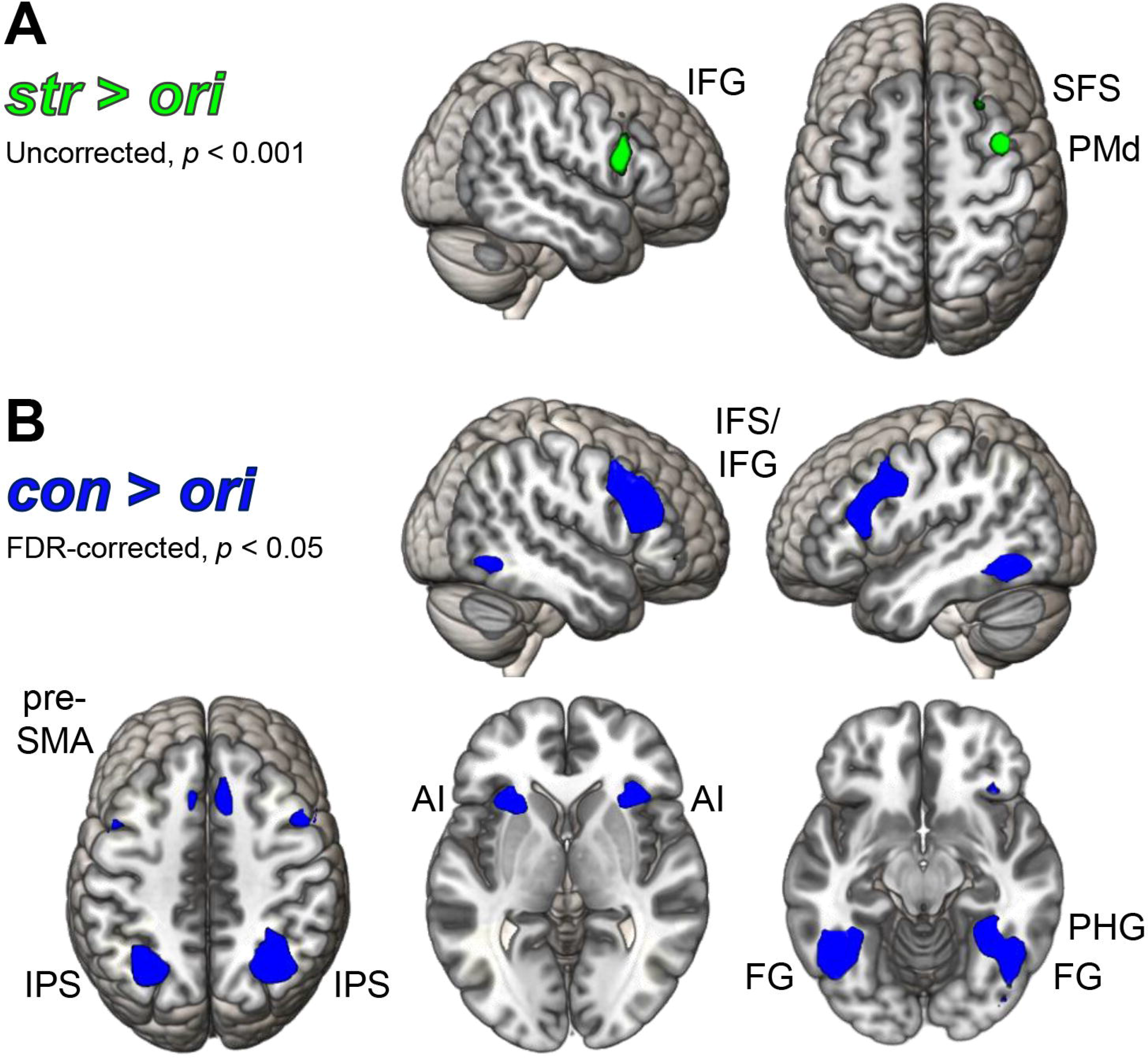
Whole brain activation for modification effects. (A) Uncorrected *t*-map (*p* < 0.001) for the *str* > *ori* contrast. (B) FDR-corrected *t*-map (*p* < 0.05) for the *con* > *ori* contrast. IFG: inferior frontal gyrus; SFS: superior frontal sulcus; PMd: dorsal premotor cortex; IFS: inferior frontal sulcus; pre-SMA: pre-supplementary motor area; IPS: intraparietal sulcus; AI: anterior insula; FG: fusiform gyrus; PHG: parahippocampal gyrus.

**Table 1.** Peak activations from second-level whole-brain analyses of modification effects.

| Localization | H | Cluster extent | MNI Coordinates |  |  | *t*-value |
| --- | --- | --- | --- | --- | --- | --- |
|  |  |  | x | y | z |  |
| *str* > *ori* (uncorrected at $p < 0.001$) | | | | | | |
| Superior frontal sulcus | R | 15 | 27 | 23 | 59 | 3.72 |
| Dorsal premotor cortex | R | 52 | 39 | 2 | 56 | 4.80 |
| Inferior frontal gyrus (pars opercularis, BA 44) | R | 76 | 54 | 14 | 20 | 4.67 |
| *con* > *ori* (FDR-corrected at $p < 0.05$) | | | | | | |
| Pre-supplementary motor area | L | 49 | -6 | 14 | 53 | 4.16 |
|  | R | 64 | 9 | 17 | 47 | 4.10 |
| Intraparietal sulcus | R | 292 | 30 | -61 | 50 | 6.23 |
|  | L | 244 | -27 | -64 | 38 | 5.47 |
| Inferior frontal junction | L | 321 | -39 | 2 | 32 | 6.43 |
| Inferior frontal sulcus, extending into inferior frontal gyrus (pars triangularis, BA 45) | L |  | -42 | 20 | 26 | 5.54 |
|  | R | 508 | 45 | 29 | 20 | 7.04 |
| Anterior insula | R | 132 | 33 | 20 | -1 | 5.57 |
|  | L | 138 | -30 | 20 | -4 | 5.68 |
| Occipitotemporal cortex | L | 239 | -45 | -55 | -13 | 7.55 |
| Fusiform gyrus | L |  | -33 | -46 | -16 | 4.39 |
| Occipitotemporal cortex, including fusiform gyrus and extending into parahippocampal gyrus | R | 266 | 33 | -52 | -13 | 5.12 |

Note: H = Hemisphere, MNI = Montreal Neurological Institute, L = Left, R = Right, *str* = structure modification, *con* = content modification, *ori* = original.

To detect brain regions which respond to content modifications, we calculated the whole brain contrast *con > ori*. Significant bilateral activation was found in pre-SMA and the middle IPS. Additionally, there were activation clusters in left and right inferior frontal sulcus (IFS) comprising the triangular part of the IFG (BA 45) and the inferior frontal junction (IFJ). Further, there was significant bilateral activation in the anterior insula and in the OTC, including bilateral FG and extending into parahippocampal gyrus (PHG) in the right hemisphere (Figure 5 and Table 1).

#### 3.3.2 ROI analysis of BA 44 and BA 45

The rmANOVA with the factors HEMISPHERE (*left*, *right*), ROI (*BA 44*, *BA 45*), and CONTRAST (*str > ori*, *con > ori*) revealed a significant main effect of HEMISPHERE (*F*_(1,35)_ = 18.458, *p* < 0.001, η*p²* = 0.345), driven by higher beta values in the right (*M_right_* = 0.359, *SE_right_* = 0.092) than in the left hemisphere (*M_left_* = 0.096 , *SE_left_* = 0.060). Further, we found a significant main effect of CONTRAST (*F*_(1,35)_ = 11.317, *p* = 0.002, η*p²* = 0.244), as beta values extracted from the *con* > *ori* contrast (*M_con_* = 0.331, *SE_con_* = 0.082) were generally higher as those extracted from the *str* > *ori* contrast (*M_str_* = 0.124, *SE_str_* = 0.073). While we found no significant effect of ROI (*F*_(1,35)_ = 0.027, *p* = 0.871, η*p²* = 0.001), there was a significant interaction of the factors ROI and CONTRAST (*F*_(1,35)_ = 50.405, *p* < 0.001, η*p²* = 0.590). Post-hoc paired *t*-tests revealed that beta values were significantly lower in BA 45 for structure (*M_str_45_* = 0.074, *SE_str_45_* = 0.075) than for content modifications (*M_con_45_* = 0.387, *SE_con_45_* = 0.084; *t*_(35)_ = -4.963, *p* < 0.001). Further, we found that beta values for structure modifications were significantly higher in BA 44 (*M_str_44_* = 0.173, *SE_str_44_* = 0.078) than in BA 45 (*t*_(35)_ = 2.349, *p* = 0.049), while the opposite was the case for content modifications (*M_con_44_* = 0.275, *SE_con_44_* = 0.086; *t*_(35)_ = -2.572, *p* = 0.029).

## 4 DISCUSSION

Episodic memories allow us to mentally relive our personal past. Against what we might believe, they are not exact imprints of previous experiences, as they are prone to later modification, potentially triggered by prediction errors. In the present study, we investigated brain and behavioral responses to retrieval cue induced prediction errors based on two properties which characterize episodic memories, namely structure and content information. While brain responses were largely as expected for content prediction errors, only subtle activation was detected following structure prediction errors. A post-fMRI memory test revealed an increased tendency to accept modified episodes when the same had been presented during the fMRI scan. Together, findings provide first evidence that different types of mnemonic prediction errors are processed differently by the brain and may contribute to memory changes.

### 4.1 Influence of episodic modifications on post-fMRI memory performance

As expected, episodes that had been presented in a modified version during the fMRI experiment were generally remembered worse, as reflected by lower discrimination indexes (*P_r_*). More specifically, modified videos were more often perceived as original stories thereafter, as evidenced by significantly more acceptive ratings and increased false alarm rates. Others have reported that mnemonic prediction errors lead to an intrusion of new information into an established memory repertoire (Long et al., 2016; Sinclair & Barense, 2018).

Previous studies report that the inclusion of new information into an old memory is not necessarily accompanied by a deterioration of the original memory (St. Jacques et al., 2013; Sinclair & Barense, 2018). Also in our study, the acceptance of original videos was only descriptively reduced after encounters of alternative versions, while hit rates indicating detection of original episodes were at ceiling. In contrast, Kim et al., showed that prediction violations led to decreased recognition of original memory content. In their study, original memories were encoded by showing the to-be-predicted sequence only once (Kim et al., 2014), whereas participants in our study had experienced and actively encoded each story multiple times. Thus, it seems plausible that memory traces were particularly stable in the current paradigm, such that a greater number of repeated alternative evidences would have been required to significantly reduce memory performance for the initially encoded episodes.

Participants took longer to correctly classify unmodified videos when they had previously experienced viewed structurally modified videos. Longer response times in cued-memory paradigms are interpreted as indicative of increased difficulty of retrieval due to higher cognitive processing demands (Larsen & Plunkett, 1987; Noppeney & Price, 2004). Thus, it is likely that it became more difficult to differentiate between alternative competing versions of episodes when versions diverging from the original experience had been already encountered in the fMRI session.

Taken together, the behavioral findings suggest that structure as well as content modifications during cueing of episodic retrieval influenced subsequent memory for these episodes. Our findings corroborate the observation that mnemonic prediction errors can trigger episodic memory modification (Sinclair & Barense, 2019). While changes in memory may sometimes reflect the updating of the original memory trace (Sinclair & Barense, 2018; 2019), our findings suggest that original memory traces remained largely intact as original episodes were still correctly recognized. Our results rather indicate that unexpected new information led to the storing of an additional, alternative version of the original episode.

What remains unclear is how exactly memory traces were influenced by our intervention. For example, it has been discussed that memory modification can result from an interference of old and new memory traces (Sederberg et al., 2011; Klingmüller et al., 2017; Sinclair & Barense, 2018; 2019) or from source confusion (Hekkanen & McEvoy, 2002), which could both explain our findings. On the other hand, participants remembered correctly that untrained videos had not been part of the original episode repertoire even though they had repeatedly been presented during the fMRI experiment as well. This finding suggests that our post-fMRI memory test results are probably not a product of source confusion in its simplest form.

Another interesting behavioral finding was that participants generally had a strong tendency to accept structurally modified versions as originals in the memory test. Although both types of modifications resulted from a single change in the story (cf. Supplementary Table 1), it is likely that structural modifications were generally less salient than content modifications. This would be matched by the fact that after the fMRI session, two-thirds of the participants reported noticing at least one object swap, while less than a quarter of them had noticed a change in the sequence of action steps. Recently, it was reported that memory performance following prediction errors differed depending on whether changes were detected (and remembered) by participants or not (Wahlheim & Zacks, 2019). While undetected changes led to reduced memory performance, detected changes had the opposite effect. Depending on contextual factors, prediction errors can even improve subsequent memory (Smith et al., 2013; Greve et al., 2017). Although our results suggest that structural changes were rarely noticed, more frequent detection in the case of content modification did not lead to enhanced, but on the contrary, also to worse memory performance. Thus, it appears that the effect of prediction violation on memory is subject to complex dynamics. Specifically, it has been put forward that strong prediction errors lead to better memory by promoting learning of new associations (Greve et al., 2017). Since weak prediction errors were also found to enhance memory performance (Greve et al., 2019), it may turn out that the strength of prediction errors need to overcome a certain threshold to initiate memory modification (Exton-McGuinness et al., 2015) or modulates memory even in a U-shaped fashion (Greve et al., 2019). The effect of the quality of prediction errors in episodic retrieval and the effect of their quantity need to be investigated simultaneously in future studies.

### 4.2 Neural responses to different episodic modifications

New content and structure information of the episodic cue was expected to draw on distinct brain areas, but also to share some common activation in MFC, areas involved in action observation, and the hippocampal formation. However, only without correction for multiple comparisons, common activation was detected in right IFG, which reflects our previous finding that IFG increases for inconsistent or highly informative detail in observed actions (Wurm & Schubotz, 2012; Hrka et al., 2015; El-Sourani et al., 2019). Here, it is not possible to distinguish whether nonsignificance ^ć^was due to a large functional disparity between content and structural information in episodic memories, or simply to an overly subtle brain response to violated structural predictions.

On the one hand, we had expected that structure modifications lead to activation in brain regions involved in the temporal organization of episodes. We did not find significant activation patterns after correction for multiple comparisons. Without such correction, we detected activation in a hypothesized area, right PMd/SFS. The activity of a region comprising PMd/SFS is related to linking successive action steps (Stadler et al., 2011; Schubotz et al., 2012; Hrka et al., 2014; Pomp et al., 2021), and could contribute to updating the current event or action mode^ć^l with respect to each next segment (Kurby & Zacks, 2008; Tamber-Rosenau et al., 2011; Schubotz et al., 2012; Pomp et al., 2021).

On the other hand, we found content modifications to specifically recruit pIPS and OTC, including FG, which were hypothesized on the basis of their role for processing of object properties in the context of actions (Grill-Spector et al., 2001; Reber et al., 2005; Wiggett & Downing, 2011; Lingnau & Downing, 2015; El-Sourani et al., 2019). More specifically, pIPS encodes basic visual features of graspable objects (Creem-Regehr, 2009; Mruczek et al., 2013), reflecting the interaction with toy objects in our paradigm. Additionally, content modifications elicited activity in the hippocampal formation (right PHG). In general, hippocampus and PHG are important in learning contexts (Aguirre et al., 1996; O’Reilly & Rudy, 2000; Davachi & Wagner, 2002; Köhler et al., 2002) and there is evidence that the PHG is involved in processing of competing memories (Kuhl et al., 2012). Moreover, the hippocampal formation is believed to generate mismatch signals when predictions do not fit perceptual inputs (Kumaran & Maguire, 2007; Duncan et al., 2009; Long et al., 2016). Because our post-fMRI memory test data imply that content changes were more salient than structural changes, one could speculate that the discrepancy between prediction and perception was strong enough to be reflected in the activation of the hippocampal formation for content modifications. This would also fit with our finding that content modifications triggered activity in the anterior insula, which is relevant for error detection (Klein et al., 2013) and salience processing (Menon & Uddin, 2010).

Consistent with our conjunction analysis, the ROI analysis showed stronger activity in the right hemisphere in different IFG regions. Activation in BA 45 was stronger for content changes than for structure changes. These results confirm the assumption that BA 45 is more responsive to content than to structural prediction errors (Badre & Wagner, 2007; Heim et al., 2009). We take this as a first indication that within IFG, different subregions might process different types of episodic prediction errors. However, our study cannot rule out the possibility that this effect is merely due to quantitative (rather than qualitative) differences between structural and content prediction errors, as structure modifications might have elicited weaker prediction errors in our paradigm.

### 4.3 Limitations and implications for future research

One factor that may limit the generalizability of our findings is that, for practical reasons, only women participated in the study. However, because the processing of episodic memory in the brain seems to be broadly similar between women and men (Nyberg et al., 2000), we are confident that our findings are applicable to a more general population. In the future, our paradigm could be adapted to circumvent such practical limitations, for example, by applying virtual reality techniques so that encoding could be detached from the true physical appearance of participants’ hands.

Second, we used new content and structure information which contrasted details of the encoded episodes to elicit prediction errors. Even though these interventions can also be interpreted as contextual and associative novelty, respectively, the unexpected new input within the familiar context will give rise to mnemonic prediction errors according to the predictive coding framework (see Reichardt et al., 2020 for a review). It would be valuable to find a way to keep novelty constant and have participants make active predictions which are then either violated or not.

Finally, another limitation of the current study is that brain responses to structure modifications were much weaker than those to content modifications, which may be explained by lower salience. This issue will be testable if stronger structural breaches of expectation are employed in episodic retrieval.

## 5 CONCLUSION

When recalling episodes, our memory can change, for instance due to prediction errors in the re- activation process. Our results suggest that structural and content prediction errors in episode retrieval differ in their neural processing. The tendency to falsely classify modified episodes as originally experienced ones increased after experiencing repeated structural as well as content-related prediction errors. Accordingly, different types of prediction errors can confuse episodic memory and possibly lead to the emergence of alternative versions of one and the same memory trace. Our results and the new paradigm introduced here may provide a fruitful starting point for further research on the mutability of our episodic memory.

## AUTHOR CONTRIBUTIONS

SS, BJ, NE and RS contributed to conception and design of the study. SS and BJ collected the data. SS performed the statistical analysis. IT provided substantial help with the data analysis. SS wrote the first draft of the manuscript. RS wrote sections of the manuscript. SS created the figures. SC, OW and RS contributed with scientific support. All authors contributed to manuscript revision, read, and approved the submitted version.

## ACKNOWLEDGMENTS

The authors thank Monika Mertens for her help during data collection. Furthermore, we thank Annika Garlichs, Christin Schwarzer, Helena Sydlik and Yuyi Xu for their assistance during the creation of stimulus material, data collection and data management. Lastly, we thank Jennifer Pomp, Lena Schliephake, Falko Mecklenbrauck and Nina Heins for advice regarding data analysis and the members of research unit FOR 2812 for valuable discussions. A preprint of this article has been posted on bioRxiv (Siestrup et al., 2021).

## DATA AVAILABILITY STATEMENT

The original contributions presented in the study are publicly available. This data can be found here: https://osf.io/34knw/?view_only=3194f4ad2fa54a0fa47f4189b2aa1f77.

## ETHICS STATEMENTS

The studies involving human participants were reviewed and approved by the Ethics Committee of the Department of Psychology, University of Münster. The participants provided their written informed consent to participate in this study.

