## Supplementary Material for "What happened when? Brain and behavioral responses to modified structure and content in episodic cueing"

### 1 SUPPLEMENTARY RESULTS

For the analysis of the post-fMRI memory test, we additionally considered hit rates and false alarm rates aside from ratings for original_MT_ and modified_MT_ videos. Original_MT_ videos were targets, while modified_MT_ videos were distractors.

We computed a non-parametric rmANOVA with the factors MODIFICATION_FMRI_ (*yes*, *no*) and VERSION_MT_ (*str*, *con*) to analyze false alarm rates. There was a significant main effect of MODIFICATION_FMRI_ (*F*_(1,34)_ = 17.087, *p* < 0.001, *ηp²* = 0.334) which was driven by higher false alarm rates for *yes* (*Median_yes_* = 0.625, *IQR_yes_* = 0.344) than for *no* (*Median_no_* = 0.50, *IQR_no_* = 0.250). Additionally, there was a significant main effect of VERSION_MT_ (*F*_(1,34)_ = 108.797, *p* < 0.001, *ηp²* = 0.762), as false alarm rates were higher for *str* (*Median_str_* = 0.813, *IQR_str_* = 0.250) than for *con* (*Median_con_* = 0.250, *IQR_con_* = 0.406). There was no significant interaction of MODIFICATION_FMRI_ and VERSION_MT_ (*F*_(1,34)_ = 3.221, *p* = 0.082, *ηp²* = 0.087).

Hit rates were close to ceiling for all factorial combinations (*Median_no_str_* = 1.0, *IQR_no_str_* = 0.0; *Median_yes_str_* = 1.0, *IQR_yes_str_* = 0.125; *Median_no_con_* = 1.0, *IQR_no_con_* = 0.0; *Median_yes_con_* = 1.0, *IQR_yes_con_* = 0.125) and were therefore not further analyzed.

We compared false alarm rates for untrained_MT_ and modified_MT_ videos using the Wilcoxon singed rank test. False alarm rates for untrained_MT_ videos were at a floor level (*Median_untr_* = 0.0, *IQR_untr_* = 0.0) and significantly lower than those for modified_MT_ videos (*Z* = -5.283, *p* < 0.001; *Median_mod_* = 0.531, *IQR_mod_* = 0.281).

### 2 SUPPLEMENTARY TABLES

Supplementary Table 1: Three examples of the 24 toy stories used in the study.

| **Example 1 (“Desert”)** | | | |
| --- | --- | --- | --- |
| **Step** | **Content modification** | **Original** | **Structure modification** |
| 1 | A dead tree standing on yellow sand appears | A dead tree standing on yellow sand appears | A dead tree standing on yellow sand appears |
| 2 | A brown horse comes in, letting its head hang | A brown horse comes in, letting its head hang | A brown horse comes in, letting its head hang |
| 3 | The horse falls over *and* *dies* | The horse falls over *and* *dies* | A **grey vulture** comes in, *expecting the horse to die soon* |
| 4 | A **black wolf** comes in *attracted by the dead horse* | A **grey vulture** comes in *attracted by the dead horse* | The horse falls over *and* *dies* |
| 5 | The wolf feeds from the dead horse | The vulture feeds from the dead horse | The vulture feeds from the dead horse |
| 6 | A white skeleton remains | A white skeleton remains | A white skeleton remains |
| **Example 2 (“Vacuum cleaner”)** | | | |
| **Step** | **Content modification** | **Original** | **Structure modification** |
| 1 | A light-blue cupboard with blue books on top comes in | A light-blue cupboard with blue books on top comes in | A light-blue cupboard with blue books on top comes in |
| 2 | A green houseplant comes in | A green houseplant comes in | A green houseplant comes in |
| 3 | A **yellow folding chair** comes in | An **orange school chair** comes in | An **orange school chair** comes in |
| 4 | A black-haired woman with a blue vacuum cleaner enters the scene *to clean the room* | A black-haired woman with a blue vacuum cleaner enters the scene *to clean the room* | A black-haired woman with a blue vacuum cleaner enters the scene *to clean the room* |
| 5 | The woman vacuums near the plant | The woman vacuums near the plant | The woman vacuums near the chair |
| 6 | The woman vacuums near the chair | The woman vacuums near the chair | The woman vacuums near the plant |
| 7 | A green snake comes in | A green snake comes in | A green snake comes in |
| 8 | The woman hits the snake with the vacuum pipe *to defeat it* | The woman hits the snake with the vacuum pipe *to defeat it* | The woman hits the snake with the vacuum pipe *to defeat it* |
| 9 | The woman flees the room | The woman flees the room | The woman flees the room |
| **Example 3 (“Mountain hike”)** | | | |
| **Step** | **Content modification** | **Original** | **Structure modification** |
| 1 | A woman (*hiker*) wearing a light-brown hat comes in, holding a wooden walking stick and a tuft of grass in her hands | A woman (*hiker*) wearing a light-brown hat comes in, holding a wooden walking stick and a tuft of grass in her hands | A woman (*hiker*) wearing a light-brown hat comes in, holding a wooden walking stick and a tuft of grass in her hands |
| 2 | A dark-grey rocky mountain range appears | A dark-grey rocky mountain range appears | A dark-grey rocky mountain range appears |
| 3 | A **brown fawn** stands on one of the rocks | A **black goat** stands on one of the rocks | A **black goat** stands on one of the rocks |
| 4 | The woman raises her hand holding the grass *to lure the fawn* | The woman raises her hand holding the grass *to lure the goat* | The goat approaches the woman *because it is trusting* |
| 5 | The fawn approaches the woman *to feed from the grass* | The goat approaches the woman *to feed from the grass* | The woman raises her hand holding the grass *to feed the goat* |
| 6 | A brown baby goat comes in *to feed from the grass as well* | A brown baby goat comes in *to feed from the grass as well* | A brown baby goat comes in *to feed from the grass as well* |
| 7 | The woman turns towards the baby goat *to let it feed* | The woman turns towards the baby goat *to let it feed* | The woman turns towards the baby goat *to let it feed* |

*Note:* Each story existed in three different versions: an original (middle column) which was encoded during training, a version with a content modification (left column) which included a different object (underlined and bold), and a version with a structure modification (right column) in which two adjacent action steps were switched compared to the original (shaded in grey). Parts written in cursive represent interpretations of the visible actions.

### 3 SUPPLEMENTARY FIGURES

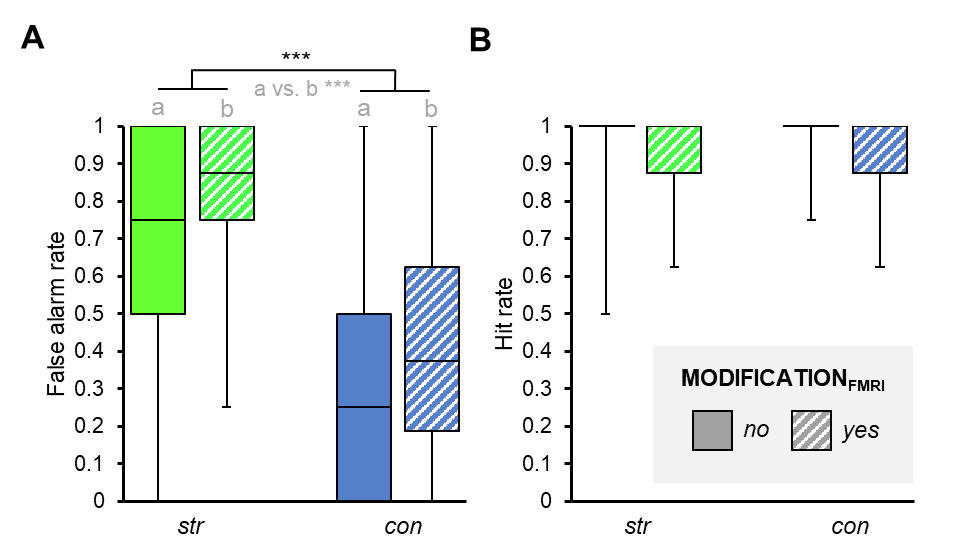

**Supplementary Figure 1:** **Supplementary results from post-fMRI memory test.** Original_MT_ videos were the targets while modified_MT_ videos were distractors. **(A)** False alarm rates. Statistics: non-parametric rmANOVA with the factors MODIFICATION_FMRI_ (*yes, no*) and VERSION_MT_ (*str, con*), *n* = 35. **(B)** Hit rates. Statistics: no statistical analysis was performed due to ceiling effects. *** = *p* < 0.001. *str* = structure modification, *con* = content modification. a vs. b indicates the main effect of MODIFICATION_FMRI_.
